# Fractal Structure of Human and Primate Social Networks Optimizes Information Flow

**DOI:** 10.1101/2023.02.23.529431

**Authors:** B.J. West, G. Culbreth, R.I.M. Dunbar, P. Grigolini

## Abstract

Primate and human social groups exhibit a fractal structure that has a very limited range of preferred layer sizes, with groups of 5, 15, 50 and (in humans) 150 and 500 predominating. This same fractal distribution is also observed in the distribution of species mean group sizes in primates. Here we demonstrate that this preferential numbering arises because of the critical nature of dynamic self-organization within complex social networks. We calculate the size dependence of the scaling properties of complex social network models and argue that this aggregate behaviour exhibits a form of collective intelligence. Direct calculation establishes that the complexity of social networks as measured by their scaling behaviour is non-monotonic, peaking globally around 150 with a secondary peak at 500 and tertiary peaks centred on 15 and 50, thereby providing a theory-based rationale for the fractal layering of primate and human social groups.

## 1. Introduction

Human personal social networks have a characteristic size (approximately 150) with a distinctive layered structure based on a hierarchically inclusive series of layers at 5, 15 and 50 within the 150 and then continuing as an external series of layers at 500 and 1500 [1]. Counting cumulatively, these layers have a very consistent scaling ratio of ∼3 across a range of contexts including (but not limited to) ego-centric social networks [2], online social networks [3,4], cellphone communication networks [4,5], and even trading networks in stock exchanges [5]. The same motif appears in the structural organisation of hunter-gatherer societies [6], the design of leisure facilities such as caravan parks [7], the size and structure of alliances in online gaming environments [8] and the structural organisation of modern armies [9]. Moreover, the same pattern has been noted in the distribution of group sizes across primate species [10], as well as in the internal structuring of primate social groups [11,12].

The fact that this pattern seems to be so widespread in so many different social contexts suggests that it is underpinned by very general structural principles. It has been noted that social group size in primates is correlated with brain size [13,14], and that this correlates in turn with environmental threats such as predation risk [14,15]. Despite these empirical relationships, it has proven difficult to find any convincing first principles explanations that might explain why social groupings across both humans and primates have the particular structure they do, and why they should form such a specific fractal pattern.

West et al. [16] provide a *prima facie* case for viewing efficiency of information flow through networks as being crucial for the structural stability of human social groups. They used two different models of group social dynamics (a decision-making model [DMM] and a swarm intelligence model [SIM]) to generate criticality-induced intelligence as a function of social group size in humans. This analysis established computationally that the scaling of the network time series varies non-monotonically with network size. In the DMM, N individuals choose between two conflicting decisions under the influence of their neighbours [17]; in the SIM, the behaviour of a group (such as the response of a flock of birds in flight from a predator) is determined by the binary choice between continuing on-course or changing course to escape [18]. These models are derivative of a class of Ising models widely used in the study of opinion dynamics (i.e. the flow of information through networks), social physics and complexity science [9,19-21]. All these models assume that spatial or social adjacency results in neighbours converging on a common opinion or behaviour through copying, infection or some form of cultural transmission. As such, they apply to a wide variety of social and other contexts.

In both DMM and SIM models, criticality is identified as a condition for optimum information flow in complex networks. Both models turn out to have criticality in their dynamics [16], with the second moment of the global time series from these models scaling as a power law in time *t*^2*δ*^. The scaling index δ has an optimum value of approximately 2/3, peaking globally at the Dunbar Number N = 150 and falling steeply away on either side of this value. The dependence of the scaling index on network size is one signature of complexity and the calculations in [16] establish that networks of size N = 150 have optimal information transmission properties, in agreement with the principle of complexity matching (PCM) [22]. The time interval *τ* between consecutive ‘crucial events’ (CEs, or state change events) is given by the waiting-time probability density function (PDF) *ψ*(*τ*), sharing in the intermediate asymptotic regime [23] with the same inverse power law (IPL) structure as the hyperbolic PDF:

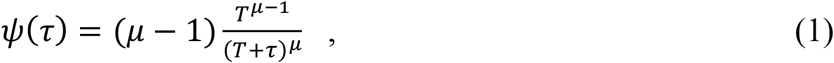

with 1 < μ < 3. The PCM has been experimentally observed in the information exchange that occurs between complex networks in a substantial number of naturally occurring interactions, including turn taking in dyadic conversation [24], the therapeutic influence of arm-in-arm walking [25], and the influence of zealots on group behaviour [17]. Following [17], the tools of network science can then be used to explore the way information flows within and between these networks in order to identify network sizes where efficiency of information flow is optimised.

Note that we use ‘information’ here in the cybernetic sense introduced by Wiener [26]. It can refer to anything that constitutes a tie or attraction between nodes in a network. In this sense, it has been used to model the economic webs of global finance and stock markets, the social meshes of governments and terrorist organizations, the transportation networks of planes and highways, the eco-webs of food networks and species diversity, the physical wicker of the Internet and the bio-net of gene regulation, as well as social relationships between individuals in a community. By the same token, although we have phrased our analysis in terms of ‘information flow’ between nodes, the results hold irrespective of what creates that flow. So, this quantity might be actual information (passed on by cultural transmission, learning or teaching) or it might be the ‘gravitational’ attraction between social partners [27] created in primates by social grooming or in humans by activities like conversation, laughter, or storytelling [28].

We here extend this approach and ask whether, in addition to the principal layer at 150 identified by [16], the other layers that have been identified in human and primate social networks [1] are also local maxima. In other words, given that the Dunbar Number of ∼150 has been shown to be the result of dynamic criticality in terms of the efficiency with which information flows through social networks [16], we use this same approach to test the hypothesis that the layers that immediately surround this value in human social groupings (i.e. network layer sizes of 15, 50 and 500) are local dynamic criticalities that represent harmonics of the Dunbar Number. If so, we may then be able to provide a first principles theoretical explanation for the network sizes that appear in both the distribution of group sizes and the fractal layering of social networks within groups – not just in humans, but also in primates.

## 2. Methods

Following [16], we consider a conventional opinion dynamics (or swarm intelligence) Ising-type diffusion model where *N* individuals (nodes) are mapped onto a *N*×*N* array. Each node is assigned an initial status synonymous with a compass direction. In successive iterations, each node can alter its status following interaction with adjacent nodes, as a result of which nodes, and the network as a whole, can switch in and out of synchrony.

We begin by relating complexity to the use of scaling theory in the search for the origin of an anomalous series ξ(t), and then use a mobile window to transform the fluctuations characterized by ξ(t) into many diffusional trajectories X(t). Stanley et al. [29,30] introduced this technique by treating DNA sequences as steps in a simple random walk (RW) process. The purpose of the RW procedure was to establish that the departure of ξ(t) from a completely random function could be detected through the departure of the scaling of X(t) from ordinary diffusion by means of a scaling index *δ* different from the default baseline of δ = 0.5. We extend this technique by interpreting X(t) as being the carrier of *crucial events* (CEs) that may be different from those hosted by ξ (t). In other words, we interpret X(t) as being a time series, which can be analysed by studying its diffusive properties. If the subsequent diffusion is anomalous, we adopt the nomenclature from the sociology literature and view the subsequent dynamics as a form of swarm intelligence model (SIM) [31].

In the case of the SIM, the fluctuations ξ(t) denote the fluctuating velocity of a swarm. It is important to stress that to detect the action of CEs, in which the time intervals between consecutive events are renewed with IPL statistics having an index μ, we convert the fluctuations ξ(t) into a diffusion process. For each of the N members of the swarm, the time step is taken for convenience to be Δ*t=*1, and the position at each successive time step is calculated using:

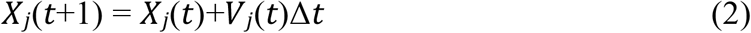

with the unit’s velocity given by a magnitude |**V**| and angle θ. The direction of the unit *j* is given by:

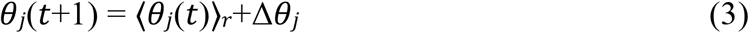

where ⟨*θ*_*j*_(*t*)⟩_*r*_ is the average direction of all units within a circle of radius *r*, at time *t* of unit *j* and Δ*θ*_***j***_ is a random number chosen from the interval [-1.75,1.75]. For every simulation the range of the random variable was the same and the constant speed had the magnitude |V| = 0.05.

The mean global field at each time step is calculated, following [18], as:

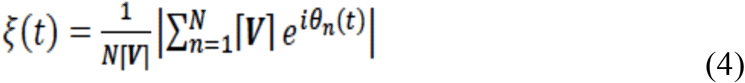

and the resulting diffusion process is:

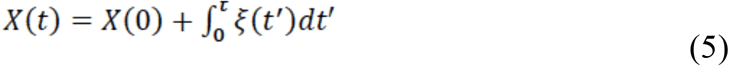

A statistical analysis of time series generated by network models of criticality-induced intelligence can then be run on the diffusion process generated by Eq. (5) for a wide range of network sizes in order to detect CEs using diffusion entropy analysis [32]. In the subsections that follow, we describe the procedure for doing so.

### 2.1. Method of stripes

We use the diffusion process generated by Eq. (5) to evaluate the scaling index δ. In the fluctuating mean field ξ(t), each fluctuation corresponding to a CE is either positive or negative and the scaling is given by δ = (μ-1)/2 for μ < 2 and by δ = 1/2 for μ > 2. This is described by the blue curve in Fig. 1. However, West et al. [33] showed that converting the negative fluctuations into positive fluctuations has the effect of making the detection of the scaling index δ much more accurate. This changing of negative to positive fluctuations corresponds to moving from the blue to the red curve in Fig. 1. It is important to stress that the results discussed herein show that the networks at criticality move from the condition δ =1/2 generated by CEs with IPL index μ = 3/2 to δ = 2/3 where the opinion persistence manifests a scaling index identical to the power law index of the crucial events [33].

**Fig. 1.**
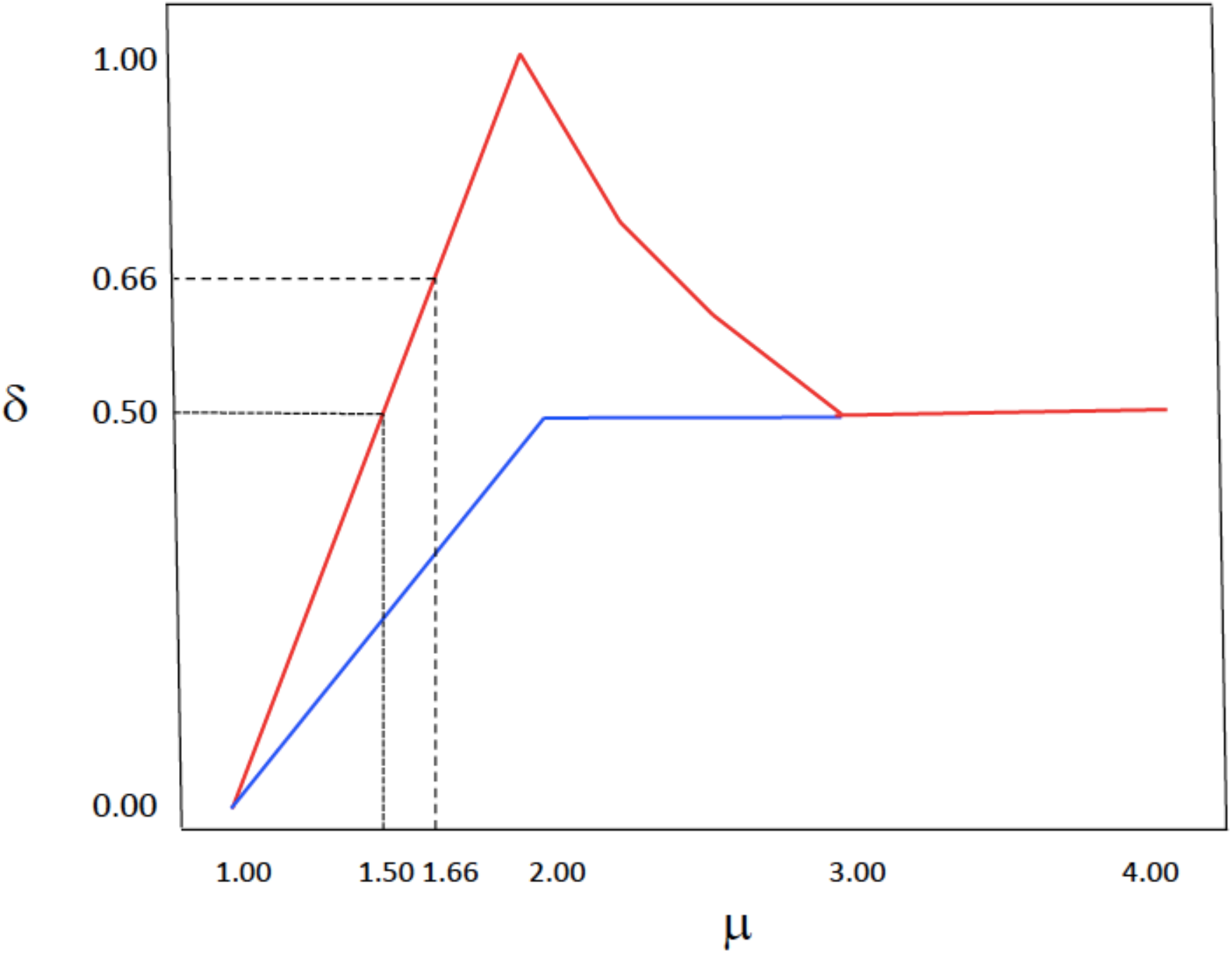
The relationships between diffusional scaling index δ and the crucial event index μ are depicted. The blue line corresponds to constraining the steps in DEA to always be positive. For this case : 1 < μ < 2, δ = (μ-1)/2. The red line corresponds to the case where this constraint is not adopted. For that case: 1 < μ < 2, δ = μ-1; 2 < μ < 3, δ = 1/(μ-1). Herein we observe values of δ in the range 0.5 ≤ δ ≤ 0.667, and values μ corresponding to them. From [33] with permission.

Following West et al. [33], the diffusion trajectory X(t) used for the scaling evaluation is built up by forcing the random walker to make a jump ahead of constant intensity when a fluctuation occurs. An alternative but equivalent approach rests on the fluctuations ξ(t) generated by SIM. Using the method of the stripes (MoS), we record the times at which the fluctuations ξ(t) cross the border between two consecutive stripes. Should these events not be crucial, they would not contribute to the scaling emerging in the long-time limit. Furthermore, when an event occurs, the random walker always makes a step ahead of constant intensity (length). The observation of the fluctuations ξ in Eq. (4) does not require the adoption of stripes but only needs the negative values of ξ to be converted into positive values.

### 2.2. Diffusion entropy analysis (DEA)

Once the diffusional trajectory is properly created on the basis of CEs, we can use a mobile window of size *l* to explore the whole diffusional trajectory of length L. For any window position, we record the difference between X(t) at the end of the window and X(t) at the beginning of the window, interpreting this as a time interval travelled by a random walker in the time interval *l*. Due to the large number of the window positions we can define the PDF *p(x,l)* and the Shannon/Wiener entropy *S(l):*

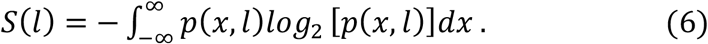

Under the assumption that the above diffusion process yields the scaling structure determined by the CEs the PDF has the scaling form:

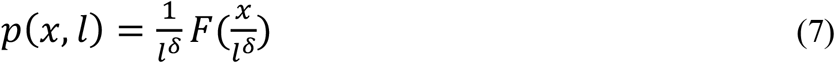

which when inserted into Eq. (6) yields:

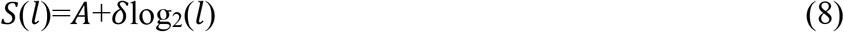

where *A* is a constant determined by the unknown function *F*(..). This reasoning enables us to interpret the slope of the curve generated by graphing *S*(*l*) versus log_2_(*l*) as the scaling index δ of the diffusion process.

### 2.3. Modified diffusion entropy analysis (MDEA)

In the present case, we use the *modified diffusion entropy analysis* (MDEA) procedure developed by [16] because it is especially suited for determining the existence of anomalies in the scaling of diffusion processes. When swarm or criticality-induced intelligence [31] becomes active, the constructed process is expected to depart from ordinary diffusion as measured by a scaling index different from δ = 0.5. Culbreth et al. [32] noted that the original version of DEA cannot assess whether the deviation from the scaling δ = 0.5 is due to the action of CEs or to the infinite memory contained in *fractional Brownian motion* (FBM) [34]. In contrast, the MDEA procedure filters out the scaling behavior of infinite stationary memory of FBM, when it exists, and the remaining departure of the scaling index from δ = 0.5 is then solely a consequence of CEs.

It is important to notice that in the DEA the diffusion trajectory X(t) is realized using the experimental data ξ(t) directly. In the more refined version given by Culbreth et al. [32] using the method of the stripes, we obtain the MDEA. Fig. 2 shows the result of this procedure in the case where MDEA is used with the stochastic equation Eq. (5). Similar results are obtained by using the detection of CEs through the stripes. It is important to stress that the adoption of MDEA makes *S*(*l*) a linear function of ln(*l*) in an extended time *l* interval. We refer to this scaling domain as the intermediate asymptotic region. At small values of *l*, the IPL index of the crucial events μ is not yet perceived. In the long-time *l* region, the deviation from the linear behavior is usually due the small number of CEs and consequently to statistical inaccuracy.

**Fig. 2.**
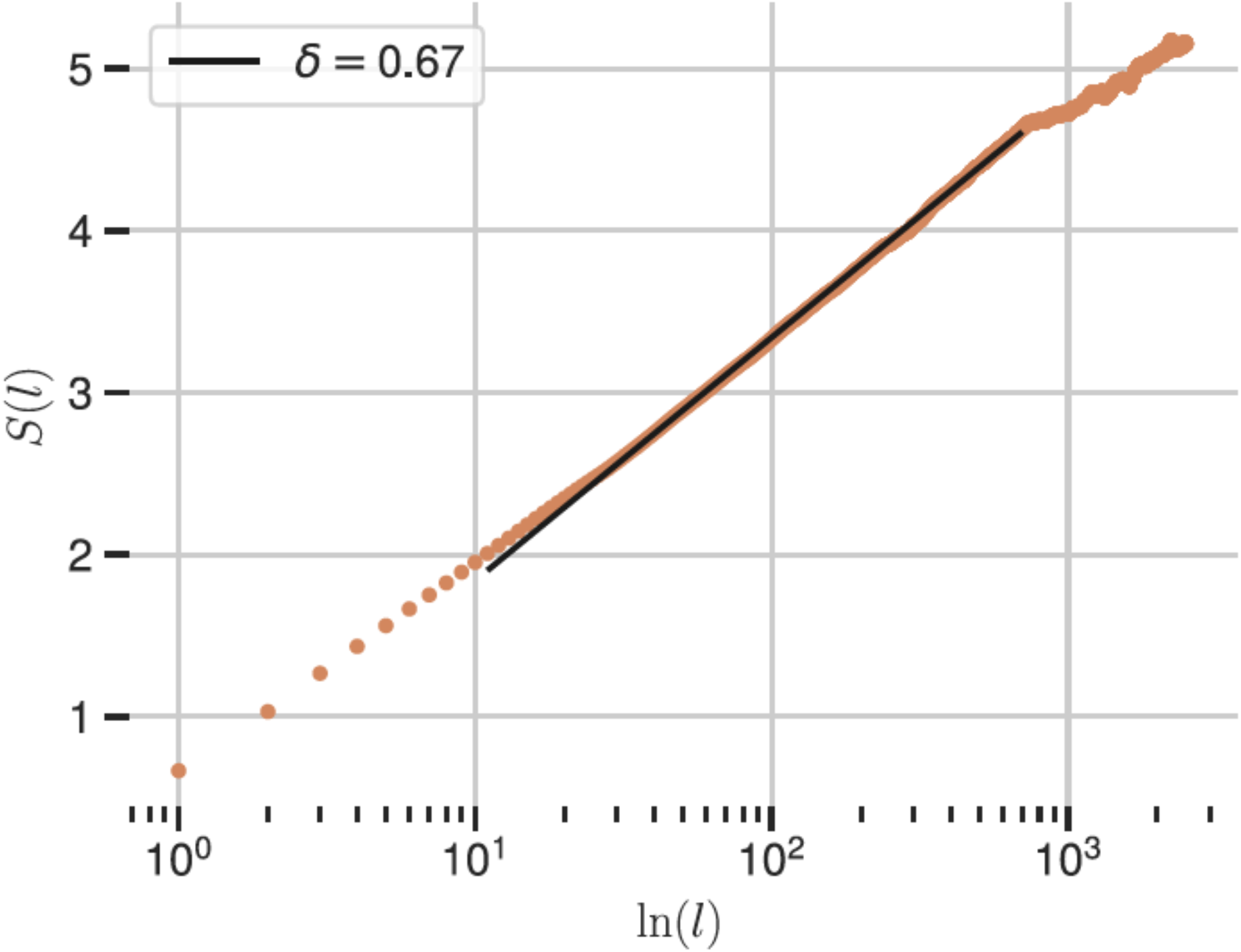
The Shannon/Wiener entropies (orange dots) at each window length *l*, measured by MDEA on time series data generated by the SIM mean field for 150- and 10,000-time steps. The scaling is the slope of the linear portion and gives δ = 0.67 (solid black line).

The MDEA method applied to the signal X(t) generated by the criticality-induced intelligence time series driving the rate equation 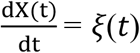 implements the original DEA in conjunction with the method of stripes to detect information regarding the phenomenon of opinion persistence. It is well known that a diffusion trajectory generated by totally random fluctuations yields a rare recursion to the origin, with the time interval between consecutive origin crossings described by the hyperbolic PDF given by Eq. (1) with μ = 1.5 [16]. Due to the forward stepping constraint on the RW in MDEA, we have: *δ* = *μ*_*S*_−1 for 1 < *μ*_*S*_ < 2 and *δ*=1/(*μ*_*S*_−1) for 2 < *μ*_*S*_ < 3, as well as δ = 0.5 for μ_*S*_≥3 [35], as shown in Fig. 3.

**Fig. 3.**
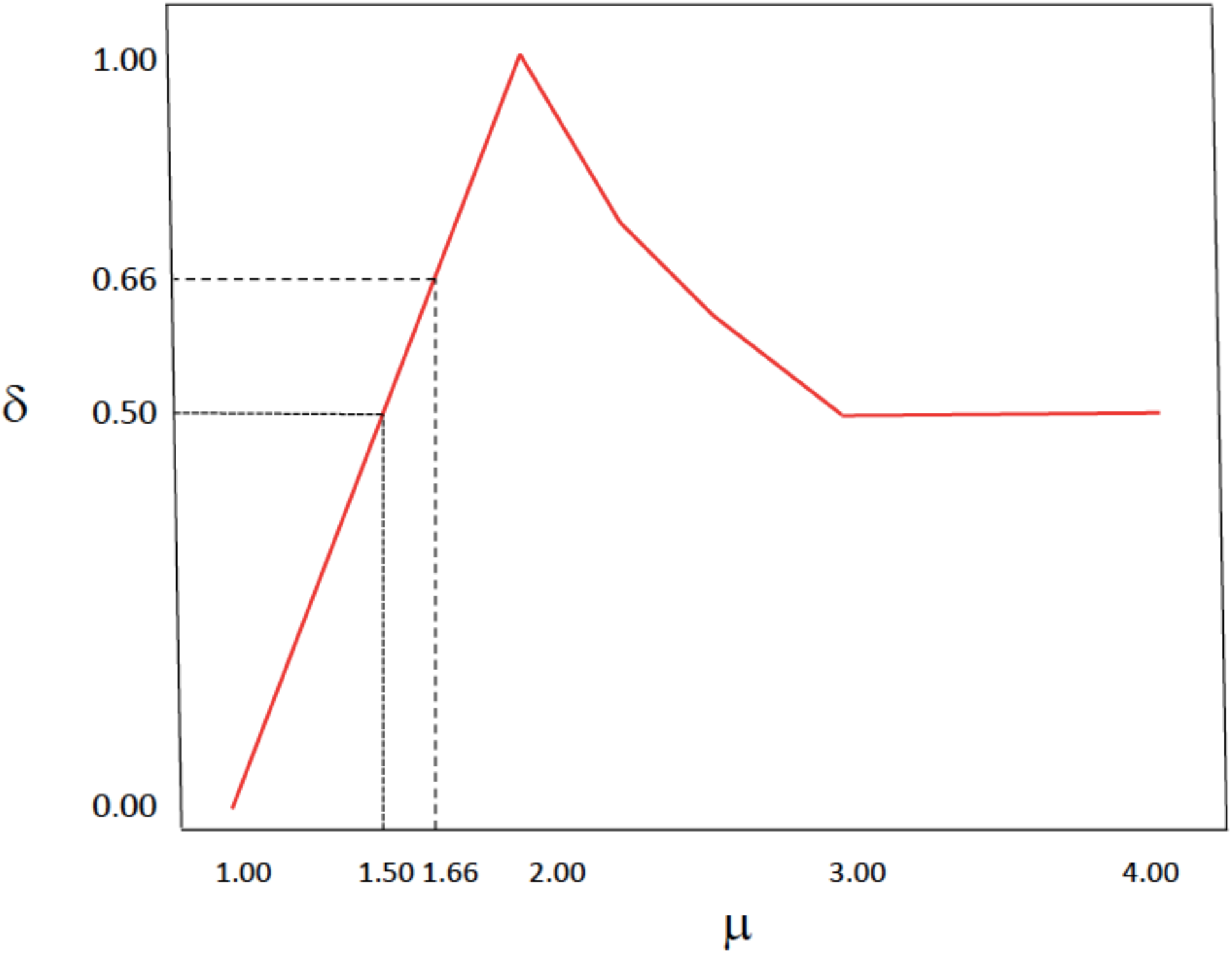
Plot of the relationship between the scaling index δ and the CE IPL index detected by using MDEA, with *μ*=*μ*_*S*_. Note that δ = 0.5 can also occur for *μ*_*S*_=1.5 and *μ*_*S*_≥3. From [36] with permission.

Notice that the value *δ* = 0.66 is the maximum scaling value occurring at N = 150. In the MDEA approach adopted herein, this scaling value signals the occurrence of the Kardar-Parisi-Zhang scaling [37]. According to the rule depicted by the solid black curve in Fig. 3, the scaling index δ = 0.5 is generated by the temporal complexity IPL index *μ*_*S*_ = 1.5 for *μ*_*S*_ < 2 and for all *μ*_*S*_≥ 3. The second condition is equivalent to an ordinary Poisson process. In summary, δ = 0.5 may be determined by both CEs (with *μ*_*S*_ = 1.5) and non-CEs (when *μ*_*S*_≥ 3).

The time interval between consecutive origin-crossings provides information about the system maintaining its state, while MDEA applied to ξ(t) detects the IPL index of CEs. Therefore, it is convenient to use the symbol *μ*_*R*_ to denote the complexity of opinion persistence (recrossing) index and the symbol *μ*_*S*_ is used to denote the index for the temporal complexity of CEs. When N ≠ 150, but where the temporal complexity index is *μ*_*S*_=1.5, we expect from Eq.(6) that the complexity opinion persistence index is *μ*_*R*_ =1.75. To evaluate *μ*_*R*_, we study the diffusional variable X(t), which spends an extended time in the region X(t) > 0 (corresponding to the system selecting the “yes” state) and an extended time in the region where X(t) < 0 (corresponding to the system selecting the “no” state). This is the opinion-persistence effect, previously mentioned. Evaluating the IPL index *μ*_*R*_ is a challenging computational problem, but we can overcome this by applying the MDEA to X(t). In this latter case, the scaling δ evaluated by MDEA yields *μ*_*R*_= 1 + δ. This scaling δ is different from the scaling obtained by observing ξ(t) directly, but the value of *μ*_*R*_ should be identical to the observation of the regression to the origin of X(t).

### 2.4. Relations among scaling indices

The MDEA applied to empirical time series ξ(t) determines the IPL index *μ*_*S*_ for the time interval PDF for the transitions between the positive (negative) to negative (positive) values of ξ(t). However, when applied to the generated diffusive process X(t) determines the IPL index *μ*_*R*_ for the time interval PDF for the transitions between the positive (negative) to negative (positive) values of X(t). A relation exists between these two scaling indices and is a consequence of the scaling property of the PDF for the diffusive trajectory:

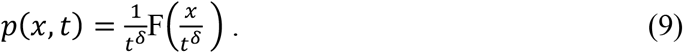

Assuming that all the trajectories of a Gibbs system are located on the origin X(0) = 0 at t = 0, we have:

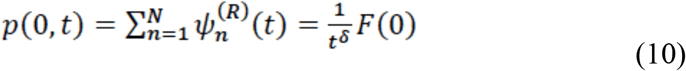

where *ψ*_*n*_^(*R*)^(*t*) is the probability that a trajectory starting from the origin at time t = 0 returns to the origin *n* times, with the last return occurring at time *t*. It is well-known that if the recrossing of the origin is a renewal process, then it is readily determined that *μ*_*R*_=2−δ. Failla et al. [38] using the CTRW establishes the relationship δ = (*μ*_*S*_-1)/2 thereby establishing the connection between *μ*_*R*_ and *μ*_*S*_:

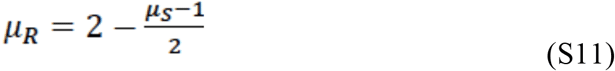

a relation proposed in [39] to account for the results obtained from the ballistic deposition model.

## 3. Results

Since the DMM and SIM models produce virtually identical results (see also [16]), we here give results for the SIM analysis only. For the SIM analysis, we hold the noise parameter (denoted as η in [18]) constant at 1.35. Agent density (the other free parameter in the model of [18]) was adjusted by holding the simulation area constant and changing the number of agents (referred to as the “number of units” in the figures), using a 2-dimensional model so that density is equivalent to the number of agents per unit area. The analysis was run across a specified set of network sizes (range 10-1000 individuals), with an average of 12.7 (range 3-28 runs per network size) (see Table 1). Since the principal focus for the analysis was on network sizes in the vicinity of the Dunbar layer sizes (15, 50, 150 and 500), the number of runs is larger in the vicinity of these values. Smaller numbers of runs were carried out for intermediate values to provide an indication of the overall pattern.

**Table 1.**
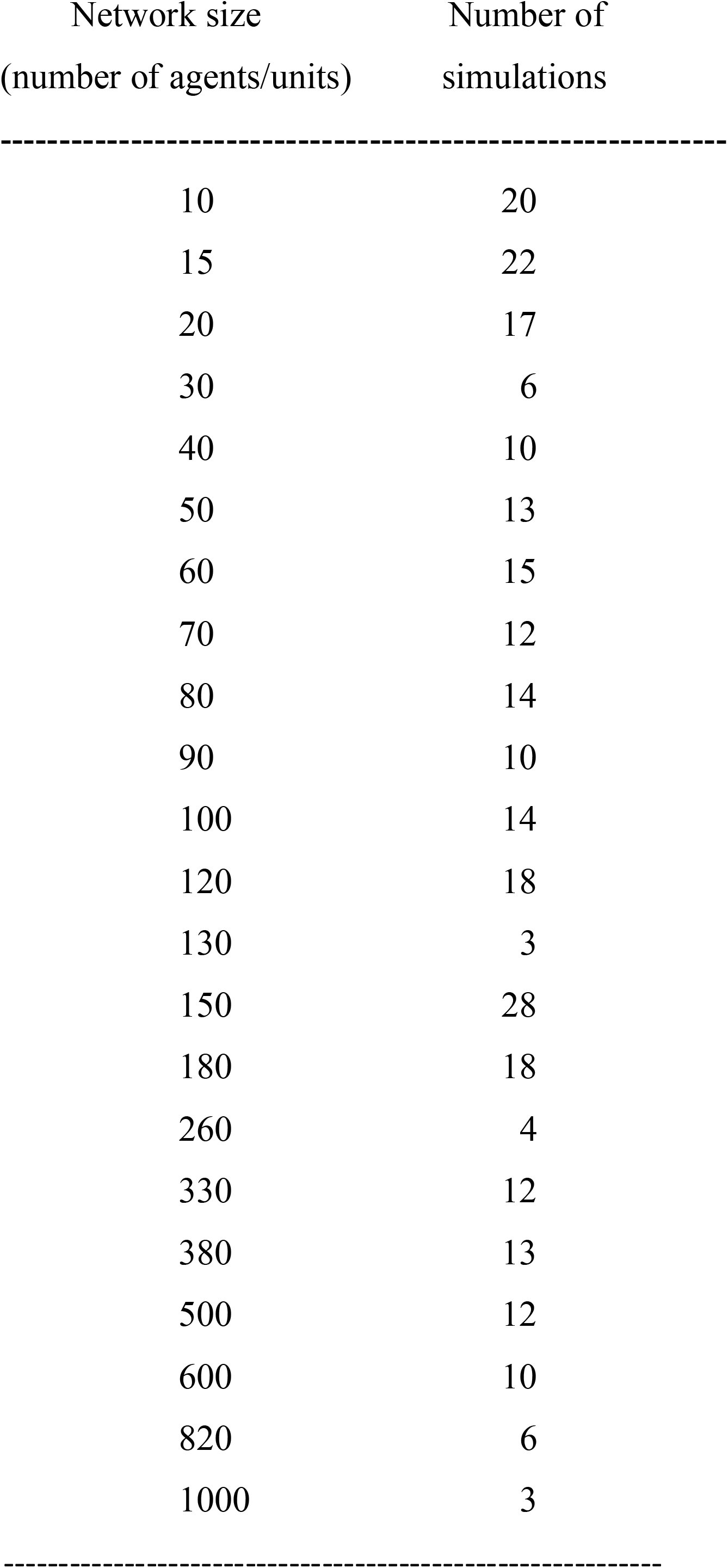
Number of runs of SIM analysis for different network sizes.

The results given below are obtained using the MDEA. This means that the fluctuation function ξ(t) of SIM is not directly used to define the crucial events. Using the method of the stripes, we record the times at which the fluctuations ξ(t) cross the border between two consecutive stripes. Should these events not be crucial, they would not contribute to the scaling emerging in the long-time limit. Furthermore, when an event occurs, the random walker always makes a step ahead of constant intensity (length). Where the time series ξ(t), with positive and negative fluctuations, is generated by CEs only, we obtain a continuous time random walk (CTRW) [36], along with the scaling, which can be properly evaluated using DEA without stripes: δ = (*μ*_*S*_−1)/2. In this case, the IPL index for CEs is given by *μ*_*R*_=2−*δ*. Under the strict condition that both CEs and opinion persistence are renewal processes we obtain, using the relation between the scaling index δ and IPL index *μ*_*R*_:

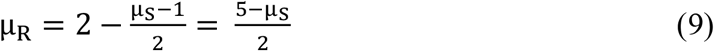

a relation originally proposed by Failla et al. [38] for studying the random growth of surfaces. We follow West et al. [16] and evaluate first the mean field for a SIM network to produce a signal. The calculations yield criticality at values of the scaling parameter whose local maximum values depend on the size of the network. Identifying the calculated value of the time rate of change of the mean field variable with the empirical time series ξ(t), we generate the RW and obtain the trajectory X(t) to which we apply the MDEA to obtain the scaling index as a function of network size N (as detailed in section 2). The resulting scaling parameter varies non-monotonically with the size of the network (Fig. 4).

**Fig. 4.**
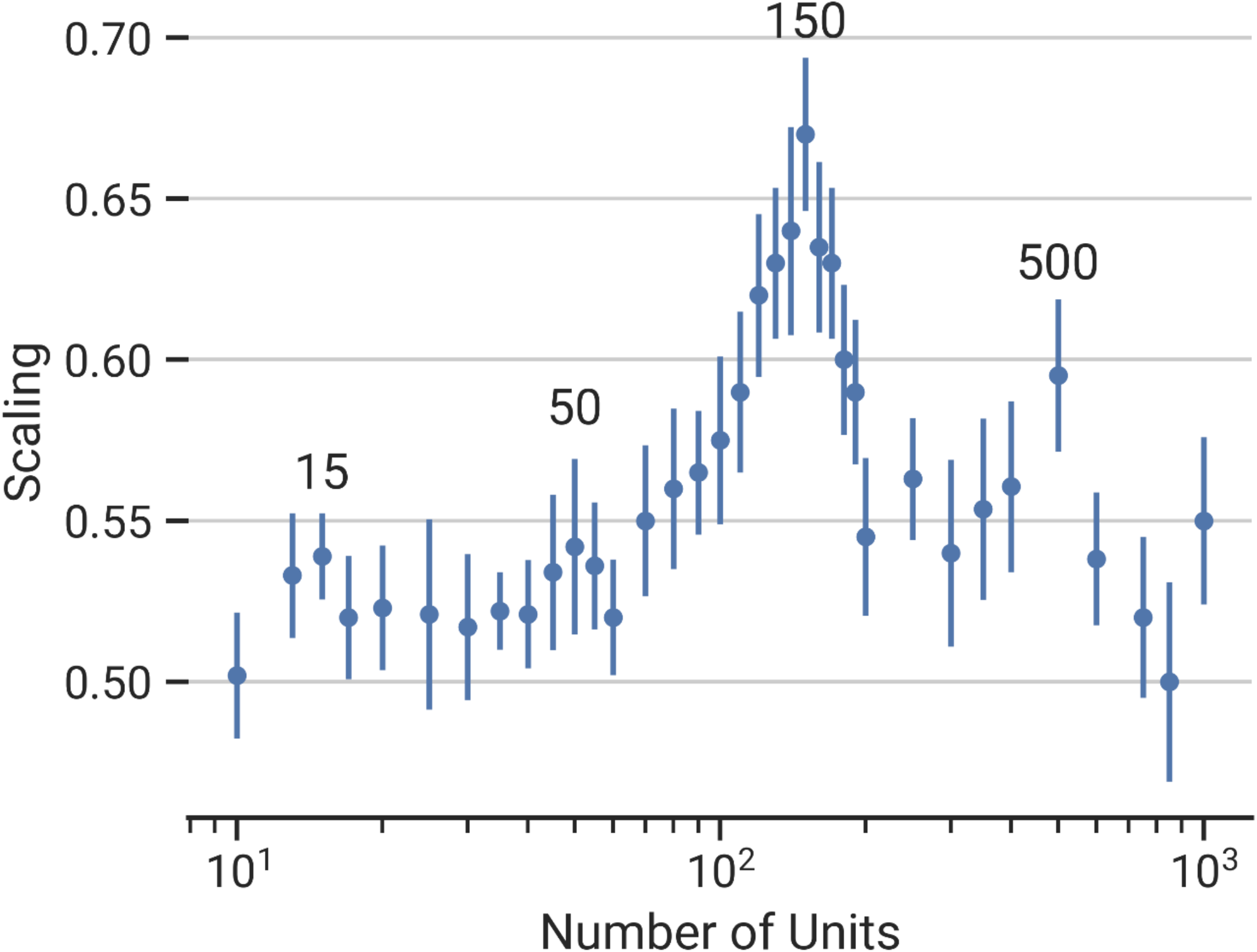
The scaling index δ shows its non-monotonic dependence on a network of size N using the SIM. Note the global maximum at the Dunbar Number 150 and the weaker secondary maximum at N = 500 and weaker still tertiary maxima at N = 15 and 50, commensurate with the layering of the Dunbar network. Symbols indicate means; error bars are standard deviations.

In order to identify peaks in the distribution of δ-values in Fig. 4, polynomials of order 2-15 were fitted to the distribution of the data in the figure. Fig. 5 plots the corresponding goodness-of-fit (indexed by the conventional r^2^) as a function of equation order. Goodness-of-fit will always increase as more terms are added to the regression equation. There are no formal methods for identifying the best fit value in these kinds of cases. However, since cumulative goodness-of-fit is invariably asymptotic, convention is to identify the value on the X-axis equivalent to the point at which the slope changes (the point of diminishing returns). On an asymptotic graph, this can be identified as the value equivalent to the point on the Y-axis that is 1/e^th^ down from the asymptote. This criterion identifies a 9^th^-order polynomial the best-fit to the data (r^2^=0.348, t_278_=12.18, p<0.0001), suggesting that there are four peaks in the data. Besides achieving a global maximum *δ*=*δ*_*m*_≈0.67 when N is at the Dunbar Number 150, the scaling index δ has additional local maxima with *δ*_*m*_>*δ*>0.5 at network sizes of N ≈ 15, 50 and 500. To confirm that the δ-values at the predicted network sizes of 15, 50, 150 and 500 are indeed local maxima, we tested whether the values of δ at each of these maxima is higher than the values immediately on either side of them. The differences in each case are statistically significant or near significant (Table 2), and, taken together as a set, the four optima are highly non-random (Fisher’s meta-analysis: p<0.0001).

**Table 2.** Comparison of δ-values in the vicinity of the predicted values network sizes of 15, 50, 150 and 500.

| Network sizes<br>peak vs comparison | t* | df | p† |
| --- | --- | --- | --- |
| 15 vs (10 + 20) | -51.87 | 52 | <0.0001 |
| 50 vs (40 + 60) | -1.51 | 33 | 0.060 |
| 150 vs (130 + 180) | -7.08 | 43 | <0.0001 |
| 500 vs (380 + 600) | -4.12 | 26 | 0.0002 |

\* unequal variances
† 1-tailed tests that the δ-values are higher on the peak than at the network sizes either side; Fisher's log-likelihood meta-analysis: $\chi^2=59.50$ , $df=2*4=8$ , $p<0.0001$ .

**Fig. 5.**
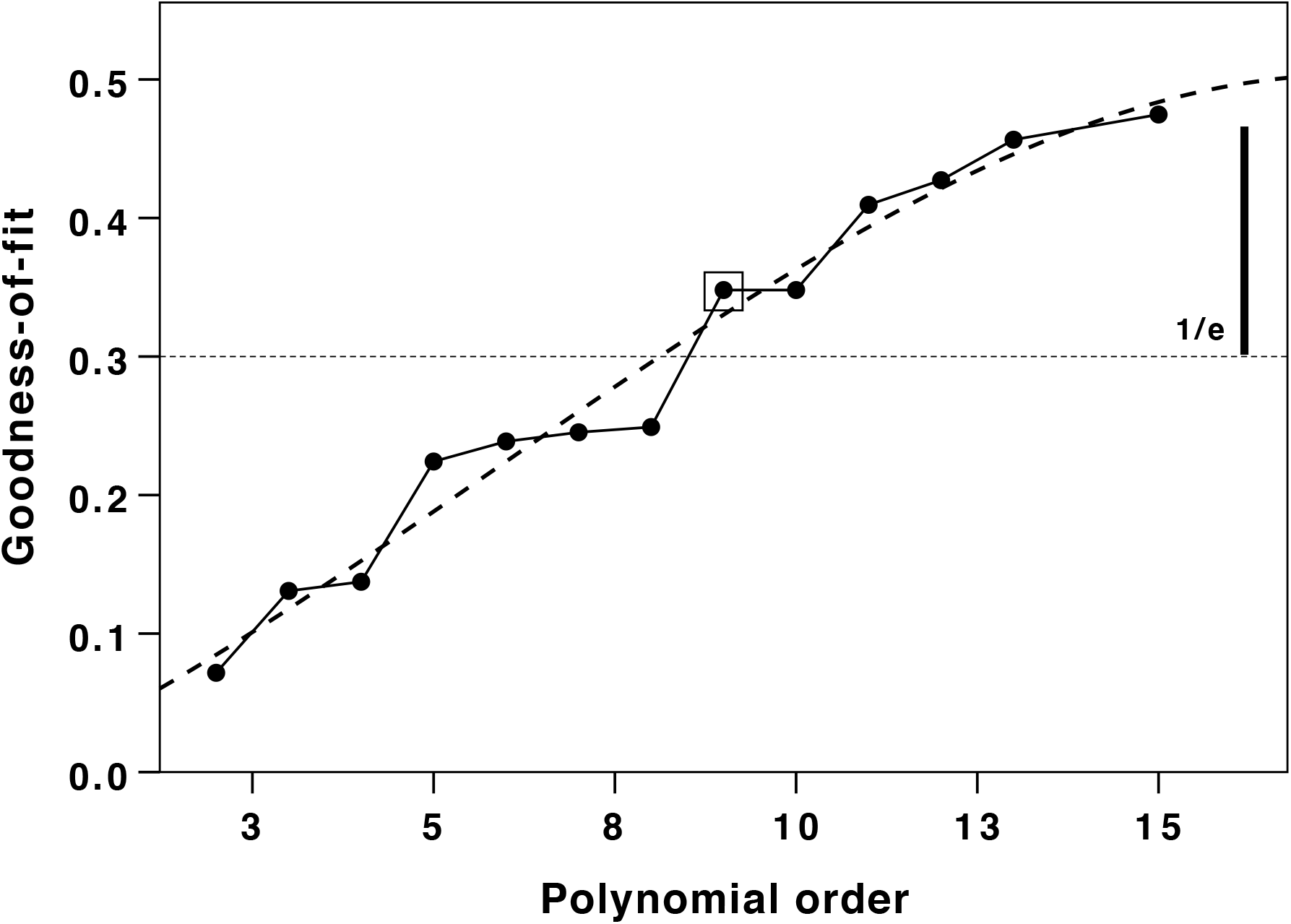
Goodness-of-fit (r^2^) for different polynomial regression fits. The long-dashed line indicates the best fit cubic regression fitted to these data. Note that goodness-of-fit approaches an asymptotic value at polynomials above 13^th^ order. The horizontal dotted line identifies the theoretical inflexion point where the slope changes to yield diminishing returns (indexed as the value on the y-axis that is 1/e^th^ down from the asymptote [indicated by thick vertical line]). The optimum polynomial order is a 9^th^ order equation (indicated by the boxed datapoint), with r^2^=0.348 (t_278_=12.18, p<0.0001). This also represents the largest absolute pairwise increase in goodness-of-fit.

We cannot emphasize too strongly that the peaks in Fig. 4 are determined only by the dynamic properties of the complex network, in the same way the single peak at the Dunbar Number was determined in [16]. Of course, just as predicting the Dunbar Number alone did not establish that this network size optimizes the transmission of information between networks but required a separate calculation, the same is true for these local maxima. This independent determination of the existence and location of layering numbers as a function of network size is made using the IPL scaling index (see [16]). The recrossing IPL index *μ*_*R*_ is determined by direct calculation from the diffusion trajectory and is compared with the theoretical values determined by the scaling parameter δ depicted in Fig. 4. The set of blue dots in Fig. 6 is obtained assuming the theoretical relation given by Eq.(9) is true and the scaling index δ has the values depicted in Fig. 6. In this case, the optima are indicated by the minima on the graph. It is evident that, despite modest divergence on the right side of the graph, the fit is excellent (comparing means: Pearson r=0.709, N=37, p<0.0001).

**Fig. 6.**
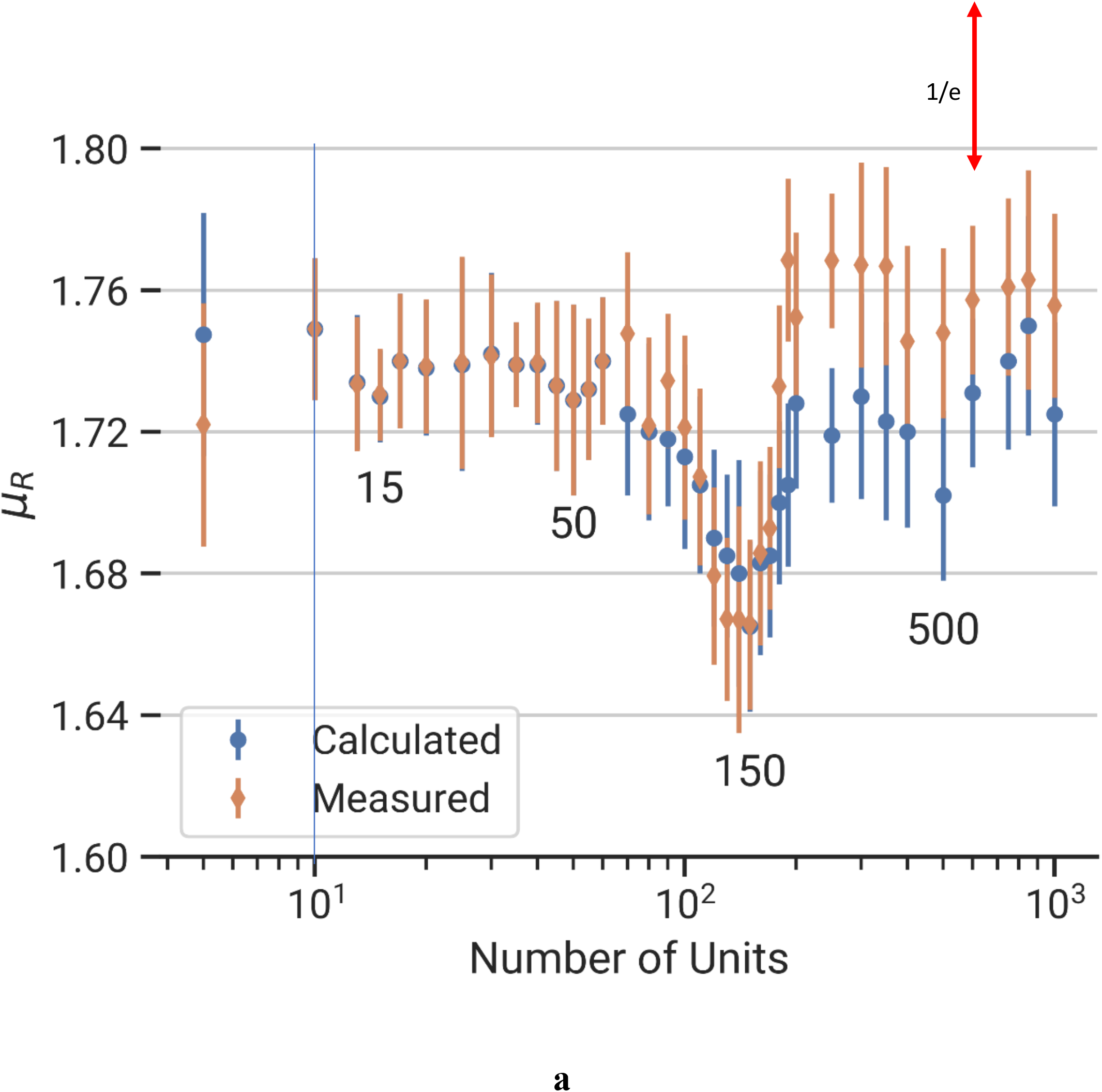
The theoretical IPL index for crucial events is given in terms of the scaling index by Eq.(2) (red points). The experimental IPL index is calculated using MDEA to process X(t) (blue points). Symbols indicate means; error bars are standard deviations.

## 4. Discussion

A substantial number of observational studies [1-8,10,11] support the concept of Dunbar layering in both human and primate social networks. Independently of the mode of communication, the numerical values of the layered structure is remarkably consistent [3,40]. The present results suggest that the distinctive sizes of these layers (as observed in the empirical data) are the product of self-organising processes in social networks that yield optimal information flow at networks of these specific values. The dependence of the scaling index on network size in Fig. 4 is one signature of complexity, and calculations in [16] present a theory-predicted value of the maximum group size for humans that agrees with the empirical Dunbar Number [1]. The theory also establishes that networks of this size have optimal information transmission properties in agreement with the principle of complexity matching (PCM) [18]. These values, thus, act as attractors both for network size and for the internal subnetwork structuring within these networks. It is of particular significance that these constraints on grouping size apply not just to humans [1] but also to the sizes and substructuring of anthropoid primate social groups as well as those of other mammals with complex multi-level social systems (e.g. elephants, orcas) [10-13,41,42].

The correspondence between theory and experiment (computation) in Fig. 6 is clearly excellent from approximately N = 10 to 200, after which point there is a 3-4% deviation between the two, with a similar deviation at the lower end (N = 5). However, a difference of this magnitude cannot be considered large and we may be requiring too much from the computation, given the simplicity of the SIM model, to expect the fit to be exact. Notwithstanding the quantitative divergence, the two sequences are highly correlated, and in Fig. 6 we observe the same qualitative dipping of the two curves in the vicinity of N = 500, with weaker dips in the vicinity of N = 15 and N = 50. Note how the values in both Figs. 4 and 6 appear to be rising towards another peak at some value of N > 1000. On the basis of the observed layering in the empirical data on human group sizes, we would expect a further peak at around N = 1500. In effect, these inner and outer optima appear to be harmonics of the central value of 150 (Dunbar’s Number).

The fractal scaling of the layers, with their distinctive scaling ratio of ∼3, seems to be a distinctive feature of human and primate social networks [1]. The fact that these layers appear to be harmonics (or subharmonics) of the Dunbar Number of 150 offers an explanation as to why these numbers appear to be unusually common in primate group sizes. It may also explain how the larger groups in the sequence are built up. There are, however, two options in this last respect. One is that when large groups are created (presumably in response to some external selection pressure), there is a natural sub-structuring of the internal network into a set of distinct sub-networks that represent local optima, producing a top-down cascade of sub-structuring to produce the layers we observe. Alternatively, groups of any given size may be produced, when required by some external need, by a bottom-up process of agglutination whereby sets of lower-level groupings are ‘bolted’ together to create a higher-level grouping: three 15-layer groups create a 50-layer, three 50-layer groups create a 150-layer, etc. If the base unit is always of a constant size in all species, this would explain why only certain group sizes are possible [10].

In real life networks, it seems that the layers in human networks, at least, are related to emotional rather than cognitive closeness [35,43,44]. This is consistent with the two-system model of the brain proposed by Kahneman [45] in which System 1 (intuition) has a fast, almost immediate response time and System 2 (cognition) has a much slower response time due to the need to organize logical thinking. It seems likely that the fast-acting intuitive part of the brain is primarily responsible for the layering structure, in support of the emotional rather than cognitive closeness of the individuals within the group, as discussed at length by West et al. [33]. The fact that the cognitive component converges on the same solution, albeit more slowly, is supported by the finding that the layers emerge naturally out of a first principles model of optimal decisions on how social capital should be invested in different alters when the benefits they offer differ in value [39,46,47].

In sum, it appears that Dunbar layering is a consequence of the nonlinear dynamics of the underlying complexity of networks that set up a series of fractally patterned attractors for group size as a consequence of efficiencies in information flow. The critical nature of the dynamics gives rise to a size-dependence of the interaction parameter thereby entailing a size-dependence on the parameter value at which criticality occurs. One way this might arise is that it is the effectiveness with which the smallest groupings are integrated that percolates through to determine the efficiency of the 150 grouping: a set of well-ordered 15-groupings necessarily creates a well-ordered 150-grouping. These results have obvious implications for our understanding of pathogen transmission, as well as the way in which social media and online multiplayer gaming environments are organised [3,8,48]. It has been suggested that several physical and chemical properties due to the thermodynamics of finite-sized-systems, including protein folding [49] and the chain length dependence of the optical properties of Perovskites [50], may analogously be due to such collective behaviour.

